# Egg size effects on nestling mass in Jackdaws *Corvus monedula*: a cross-foster experiment

**DOI:** 10.1101/2022.10.13.512039

**Authors:** Mirjam J. Borger, Christina Bauch, Jelle J. Boonekamp, Simon Verhulst

**Affiliations:** Behavioral and Physiological Ecology, Groningen Institute for Evolutionary Life Sciences, University of Groningen, P.O. Box 11103, 9700 CC, Groningen, The Netherlands; School of Biodiversity, One Health & Veterinary Medicine, University of Glasgow, Bearsden Road, G61 1QH, Glasgow, The United Kingdom

## Abstract

Variation in developmental conditions is known to affect fitness in later life, but the mechanisms underlying this relationship remain elusive. We previously found in jackdaws *Corvus monedula* that larger eggs resulted in larger nestlings up to fledging. Through a cross-foster experiment of complete clutches we tested whether this association can be attributed to egg size per se, or to more proficient parents producing larger eggs and larger nestlings, with the latter effect being more or less independent of egg size. Due to other manipulations post-hatching, we primarily investigated effects on nestling mass on day 5, which we show to predict survival until fledging. We introduce a new statistical approach to compare the competing hypotheses and conclude that 92% of the association between egg size and nestling mass is attributed to a direct effect of egg size, and that this relationship is not affected by variation in other parental quality. Intriguingly, the effect of egg size on day 5 nestling mass was steeper (1.7 g/cm^3^) than the effect of egg size on day 1 hatchling mass (0.7 g/cm^3^). Early growth is exponential, and the difference in effect size may therefore be explained by hatchlings from large eggs being further in their development at hatching. The direct effect of egg size on nestling mass raises the question what causes egg size variation in jackdaws.

## Introduction

Early-life developmental conditions can have important long-term fitness consequences (Lindström, 1999; Van de Pol et al., 2006b; Boonekamp et al., 2014). In birds, developmental conditions at the start of life are shaped by the size and content of the egg, together with its incubation. Egg size has been shown to affect offspring quality at least in early life (Krist, 2011), and sometimes also at later stages (Verhulst and Salomons, 2004; Whittingham et al., 2007; Krist, 2011; Krist and Munclinger, 2015). However, associations between egg size and offspring quality can reflect both direct effects of the egg on offspring quality as well as other parental or environmental effects after hatching when individuals in favourable environments (e.g. abundant food, few parasites) are better in raising chicks and also lay larger eggs. To resolve the question to what extent egg size directly affects offspring quality, it is necessary to experimentally separate effects of egg size from other aspects of the developmental conditions such as parental proficiency.

To this end, cross-foster experiments of complete clutches have been carried out, using different experimental designs. In non-random cross-foster experiments, clutches with large eggs were swapped with clutches with small eggs, and usually included a control treatment where large egg clutches were swapped with large egg clutches and small egg clutches with small egg clutches (Bolton, 1991; Amundsen et al., 1996; Blomqvist et al., 1997; Styrsky et al., 1999; Risch and Rohwer, 2000; Arnold et al., 2006; van de Pol et al., 2006a; Silva et al., 2007). Alternatively, clutches were swapped randomly with respect to egg size (Hipfner and Gaston, 1999; Hipfner, 2000; Bize et al., 2002; Pelayo and Clark, 2003; Krist, 2009; Reed et al., 2009). Results from these studies are mixed: Only egg effects (Amundsen et al., 1996; Hipfner and Gaston, 1999; Styrsky et al., 1999; Hipfner, 2000; Pelayo and Clark, 2003; Krist, 2009; Reed et al., 2009), only parental effects (Arnold et al., 2006; van de Pol et al., 2006a) and both egg and parental effects were found (Bolton, 1991; Blomqvist et al., 1997; Risch and Rohwer, 2000; Bize et al., 2002; Silva et al., 2007; Hadfield et al., 2013). Note that for brevity we refer to all aspects of the environment as experienced by the offspring other than the contents and size of the egg from which it hatches with ‘other parental effects’.

It would be of interest to unravel the cause of the variation in experimental results, but this is hampered firstly by the limited number of studies with varying experimental designs, but also by the analytical approach taken in most studies. Data were usually analysed by comparing whether the egg size of genetic or foster parents best explained variation in nestling survival and growth. While this approach does provide some insight, a drawback is that it does not yield a quantitative assessment of the relative contribution of egg size and other parental effects to offspring survival and growth, and that weak effects might not be detected easily. This is unfortunate, because from a naïve perspective one would expect a positive association between egg size and offspring prospects to arise through a combination of egg and other parental effects, because only when offspring prospects increase with egg size is there reason for higher quality parents to lay larger eggs. The most informative outcome of this kind of experiment is therefore an estimate of the relative contribution of the two factors to variation in offspring growth and survival. Testing whether the size of the eggs from which chicks hatched or size of the eggs produced by the foster parents best explains offspring growth does not provide information on this question; hypotheses are either rejected or not, instead of yielding an estimate of the relative importance of egg size versus other parental effects in causing the association between egg size and offspring performance. Moreover, this approach makes inferences on the basis of the absence of significant effects (e.g. no effect of original egg size on growth of offspring from cross-fostered eggs), which is a weak basis for any inference because there can be other causes for the lack of statistical significance (Milinski, 1997). To what extent egg size or other parental effects contribute to associations between egg size and offspring performance is therefore largely an open question, despite the existing studies.

We here report the results of a cross-foster experiment in a free-living population of jackdaws, *Corvus monedula*, that we performed to test to what extent a previously observed correlation between egg size and fledgling mass (Verhulst and Salomons 2004) can be attributed to a direct effect of egg size. We cross-fostered whole clutches matched for lay date and clutch size, but randomly with respect to egg size. We introduce a new approach for the statistical analysis of this type of experiment, that accepts or reject hypotheses based on significant effects rather than the absence of significance. In this approach, we considered the following hypotheses: (i) Egg size fully explains the observed correlation, in which case chicks will grow according to their own egg size, regardless of the egg size their foster parents had produced themselves (Fig. 1). (ii) Other parental effects fully explain the observed correlation, in which case chicks will grow according to the foster parent’s egg size, independent of the size of the egg from which they hatched (Fig. 1). (iii) Both egg and other parental effects contribute to chick growth, in which case chick growth will be intermediate to what is expected according to their own egg size and the size of the eggs produced by the foster parents (Fig. 1). We test these hypotheses by comparing the coefficients of the slopes represented in Figure 1, which enables us to estimate the relative contribution of both factors to chick growth.

**Figure 1:**
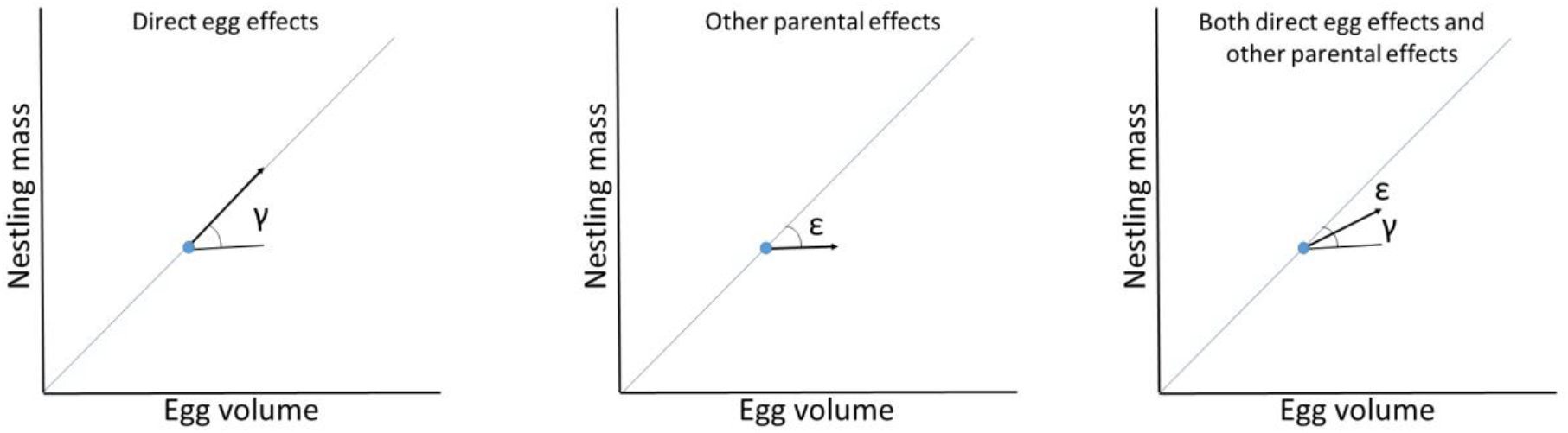
Schematic representation of the predictions following hypotheses H1 (egg size effect only), H2 (parental effect only) and H3 (egg and parental effect combined). The diagonal line represent the correlation between (average) egg volume and nestling mass on a between-individual level (i.e. this method explains differences in egg volume between individuals, no differences within the nest). The blue dots and arrows indicate the effect on nestling mass when a pair is experimentally given larger eggs than they produced themselves (as will be the case in 50% of pairs receiving a foster clutch). Following H1 (egg effect only), foster chicks will grow according to their own egg volume. In this case γ will explain the entire slope and ε will be 0. Following H2 (parental effect only) foster chicks will grow according to the egg volume of their foster parents, and hence the increase in egg volume has no effect on nestling mass (γ = 0), and ε will explain the whole slope. Following H3 (egg and parental effects combined), the slope will be explained both by γ and ε, the value of each parameter indicating which proportion of the slope is affected by egg effects (γ) and parental effects (ε). The same reasoning applies when a pair receives eggs that are smaller than they produced themselves.

## Methods

### Data collection

We studied free-living jackdaws breeding in seven colonies, each containing 9 to 22 nest boxes, south of Groningen, the Netherlands (53°08’09.9”N, 6°38’54.4”E). In our study population, jackdaws lay one clutch per year and have an average clutch size of 4.54 eggs (SD=0.97). In jackdaws there is virtually no extra-pair paternity and no intra-specific brood-parasitism (Liebers and Peter, 1998; Henderson et al., 2000), and hence the caring parents can be assumed to be the genetic parents.

From the start of the breeding season, all nests were checked once every 3 days to establish the laying date and (with limited resolution) the laying order, as jackdaws generally lay one egg per day (Verhulst and Salomons, 2004). New eggs were numbered with a felt tip pen and their length and width were measured to the nearest 0.1 mm using a sliding calliper. Egg volume (V), in cm^3^, was calculated using the formula: V = π * W^2^ * L * K /6, where W is egg width (mm), L is egg length (mm) and K = 0.00096 (Soler, 1988). Eighteen days after the first egg was laid, the nests were checked daily for newly hatched chicks. Because usually multiple eggs hatch on the same day, particularly the first eggs of the clutch, we can link only part of the hatchlings to the egg from which they hatched. Hatchlings were weighed to the nearest 0.1 gram and nail tips were clipped for individual identification. On day 5 of the brood, with day of hatching of the first hatchling counted as day 1, all chicks were weighed to the nearest 0.5 gram and unhatched eggs were removed. Parental birds were identified by their unique combination of colour rings including a numbered metal ring and unringed adults were captured during the nestling phase and ringed accordingly.

The population has been continuously studied since 2005. We cross-fostered complete clutches with the same clutch size and laying date (± 1 day) in the years 2015-2017, mostly within colonies (Table 1). In all other respects the cross-fosters were performed randomly. As a control, we used all clutches with the same range of clutch size and lay date as the cross-fostered clutches within each of the three years, i.e. clutches with a lay date before or after the first and last cross-fostered clutches of the season were omitted from the analyses, and the same selection was applied with respect to clutch size. Notably, all eggs were removed from the nest box to measure their size and hence both control and experimental clutches were treated similarly.

**Table 1:** General information about the clutches used in the data analysis of the cross-foster experiment. Numbers in brackets are the number of clutch swaps originally performed; some cross-fostered clutches were not used in the analysis due to incomplete data, usually due to the clutch not hatching.

|  | 2015 | 2016 | 2017 |
| --- | --- | --- | --- |
| *Number of swapped clutches* | 20 (20) | 37 (38) | 36 (42) |
| *Number of control clutches* | 22 | 44 | 32 |
| *Number of chicks from swapped eggs* | 78 | 138 | 111 |
| *Number of control chicks* | 82 | 145 | 110 |
| *Average egg volume (±S.D.)* | 10.81 (±1.13) | 10.93 (±1.19) | 10.50 (±1.22) |
| *Clutch size range* | 4-5 | 3-7 | 3-6 |
| *Average clutch size (±S.D.)* | 4.76 (±0.43) | 4.68 (±0.83) | 4.62 (±0.81) |
| *Lay date range (day in April)* | 15-19 | 10-18 | 8-19 |
| *Average lay date (day in April, ±S.D.)* | 16 (±1.27) | 15 (±1.93) | 13 (±3.00) |
| *Time of swapping (days after last laid egg, ±S.D.)* | 6.8 (±1.36) | 7.8 (±1.67) | 6.6 (±1.86) |

On day 5, brood size was manipulated, either enlarged or reduced by two nestlings. Two broods, one of each manipulation/treatment group, were matched based on hatch date and original clutch size. See Boonekamp et al. (2020) for further details. All fieldwork was performed by a team of researchers due to the workload, so we cannot rule out effects of differences between researchers, yet we do not think this is a big problem as all procedures were standardised.

### Data analysis

We tested which hypothesis (Fig. 1) best explained the data by estimating the slopes of the relation between nestling mass and egg volume in different models, fitted on all control and cross-fostered clutches combined. See table 2 for an overview of the models and predictions regarding the coefficients based on the different hypotheses. Note that we used this method both on nestling day 5 mass, as well as on hatchling mass. Model (1) tests whether the experimental change in egg volume (experienced by parents) affected nestling mass given the original egg volume (E_gp_), as it contains ΔE_gp-fp_, the difference in egg volume between the genetic and foster parents (arrows in Fig. 1). When γ is significantly positive, this implies that a difference in egg volume relative to the foster parents’ egg volume E_fp_ affected nestling mass, indicating a direct effect of egg volume on offspring mass. When β and γ are equal, this implies that hypothesis 1 best explains the data, i.e. there is a direct egg effect, and no evidence for an additional parental effect (see Appendix 1). This is so because E_fp_ and ΔE_gp-fp_ add up to E_gp_, the egg volume from which the chicks hatched (also for control clutches where ΔE_gp-fp_ = 0). Conversely, at the other extreme, when γ = 0, this would imply there is an effect of other parental effects only, and no direct effect of egg volume (hypothesis 2). However, non-significance of γ does not constitute strong evidence, as non-significance can have causes other than there being no effect (e.g. small sample size). Hence, we formulated model 2, which differs from model 1 in that E_fp_ is replaced with E_gp_. In model 2, δ is the dependency of nestling mass on the egg volume of the genetic parents (i.e. the volume of the eggs from which the chicks hatched) and ε is the dependency of mass on the difference in egg volume between the genetic and foster parents. Please note that the meaning of β and δ depend on the hypothesis that best fits the data, and therefore giving an unambiguous definition is not possible.

**Table 2:** Overview of hypotheses, and the models to test these. See Figure 1 for an illustration of the coefficients γ and ε, or see Appendix 1 for a mathematical explanation of all coefficients. B_a_ denote the intercepts and e_a_ are the residual variances. E_gp_ is the average egg volume of the genetic parents, E_fp_ the average egg volume of the foster parents and ΔE_gp-fp_ is the difference between these two egg volumes.

| Models |  |
| --- | --- |
| Model 1 | $\text{Mass} = B_1 + \beta * E_{fp} + \gamma * \Delta E_{gp-fp} + e_1$ |
| Model 2 | $\text{Mass} = B_2 + \delta * E_{gp} + \varepsilon * \Delta E_{gp-fp} + e_2$ |
| Hypotheses | Predicted coefficients |
| 1: Only direct egg effect | $\beta=\gamma=\delta, \varepsilon=0$ |
| 2: Only parental effect | $\beta=\delta=-\varepsilon, \gamma=0$ |
| 3: Both egg and parental effects | $0<\gamma, \varepsilon<0$ |

When there are only direct egg effects (hypothesis 1), β will be equal to γ and δ, while ε will be zero, because only the volume of the eggs from which the chick hatched determines its mass, i.e. there is no additional parental effect. Alternatively, when there are other parental effects, ε will be negative, indicating that parents receiving larger eggs then they produced themselves rear these young to a lower mass than parents would have done that produced eggs with that volume themselves. In the extreme case that there are other parental effects only, ε will equal -β (see Appendix 1). Again, this is so because E_gp_ - ΔE_gp-fp_ is equal to E_fp_, the egg volume that the foster parents produced. In the intermediate case, with both direct egg effects and other parental effects (hypothesis 3), the coefficients will take values that are intermediate to the extremes (hypotheses 1 and 2). This is where the added value of our approach emerges, because hypothesis 3 requires both γ and ε to deviate significantly from zero, and hypotheses 1 and 2 require either γ (H1) or ε (H2) to deviate significantly from zero – thus ensuring that hypotheses are in no case accepted on the basis of the absence of a significant effect. Moreover, γ / (γ + |ε|) yields an estimate of the relative contribution of direct egg volume effects to the association between egg volume and offspring mass, while its complement |ε| / (γ + |ε|) yields an estimate of the relative contribution of other parental effects.

Our hypotheses refer to between clutch variation in egg volume, and therefore in the models outlined above we used mean egg volume of the clutch throughout. However, egg volume variation within clutches may also affect nestling mass. We accounted for this in the models by including the difference between average egg volume within a clutch and an egg’s own volume in the models (van de Pol et al., 2006a; van de Pol and Wright, 2009). The accuracy with which the egg could be identified from which a chick hatched was variable, and in the analyses we used the mean egg volume of all eggs from which a chick could have hatched (e.g. if two chicks were found to have hatched since the last visit, the average egg volume of both these eggs was used as the egg volume for both chicks).

### Body mass and survival

Body mass was first measured on the day of hatching (day 1) and again for the entire brood on day 5 of the oldest chick. To account for age differences within broods, the days elapsed between hatch date of the oldest sibling and hatch date of the focal chick was included as fixed effect in the models.

To account for non-independence of measurements, genetic female ID, colony and dyad identity (linking two clutches that were cross-fostered) were also added to the model as random effects. Nest ID was not added as a random effect, since the combination of female ID and dyad ID already accounted for this variation.

We used the day 5 mass as a proxy of chick quality, since brood size manipulation experiments that involved further cross-fostering were performed on day 5 (Boonekamp et al 2020). To verify whether day 5 mass is predictive of subsequent growth and survival, and hence an informative proxy of nestling quality, we tested whether day 5 mass predicted survival until day 30, which is about 5 days before fledging. As described above for egg volume, we partitioned post-manipulation day 5 mass variation into between- and within-nests components (van de Pol et al., 2006a; van de Pol and Wright, 2009). Since broods were manipulated at day 5, we used the average and deviation of the brood average after manipulation in this model. We used a binomial distribution for the survival analysis, and included genetic female ID, Colony and Year as random effects in the model. Since this question/analysis is independent of the clutch cross-foster experiment, we included a maximum of available data (from 2005 until 2017) in this analysis.

All analyses were done in R 3.4.1 (R Core Team, 2017), with packages Lme4 (Bates et al. 2015), LmerTest (Kuznetsova et al. 2017) and MuMIn (Bartón 2019). For data visualisation ggplot2 (Wickham, 2016) and cowplot (Wilke, 2020) were used.

## Results

### Clutch swaps

Clutch swaps were performed on average 7.0 (S.D.=0.18) days after the last egg was laid, varying from 6.6 to 7.8 days between years (Table 1) and in total 100 clutches were swapped.

The average absolute difference in egg volume between the swapped clutches was 0.07 ± 1.17 cm^3^ (Fig. S1), i.e. 0.67% of the average egg volume (Table 1). From the 100 swapped clutches, seven were excluded from the analysis: three clutches because the eggs did not hatch, in three broods all chicks had died before day 5, indicating predation or parental desertion, and in one case mass was not measured on day 5. The 93 remaining swapped clutches had a total of 327 nestlings on day 5 (see table 1 for a breakdown per year) and there were 98 control clutches with 337 nestlings.

### Hatchling mass

Cross-fostering can in principle affect incubation and thereby hatchling mass, and we therefore first investigated effects of egg volume on hatchling mass (n = 648 chicks, of which 322 were cross-fostered). The results are summarized in table 3, and show that β, γ and δ were indistinguishable at 0.67 g/cm^3^, while there was no effect (0.01 g/cm^3^) of ε, indicating only egg and no other parental effects on hatchling mass (table 3). There was no effect of a clutch being cross-fostered on hatchling mass (tested by adding the factor cross-fostered (Y/N) to the model; table 3). Within clutch variation in egg volume similarly affected hatchling mass as between clutch variation indicating that also within clutches larger chicks hatched from larger eggs (slope of 0.76 ± 0.16 g/cm^3^; p<0.001, table 3, Fig. 3.

**Table 3:**
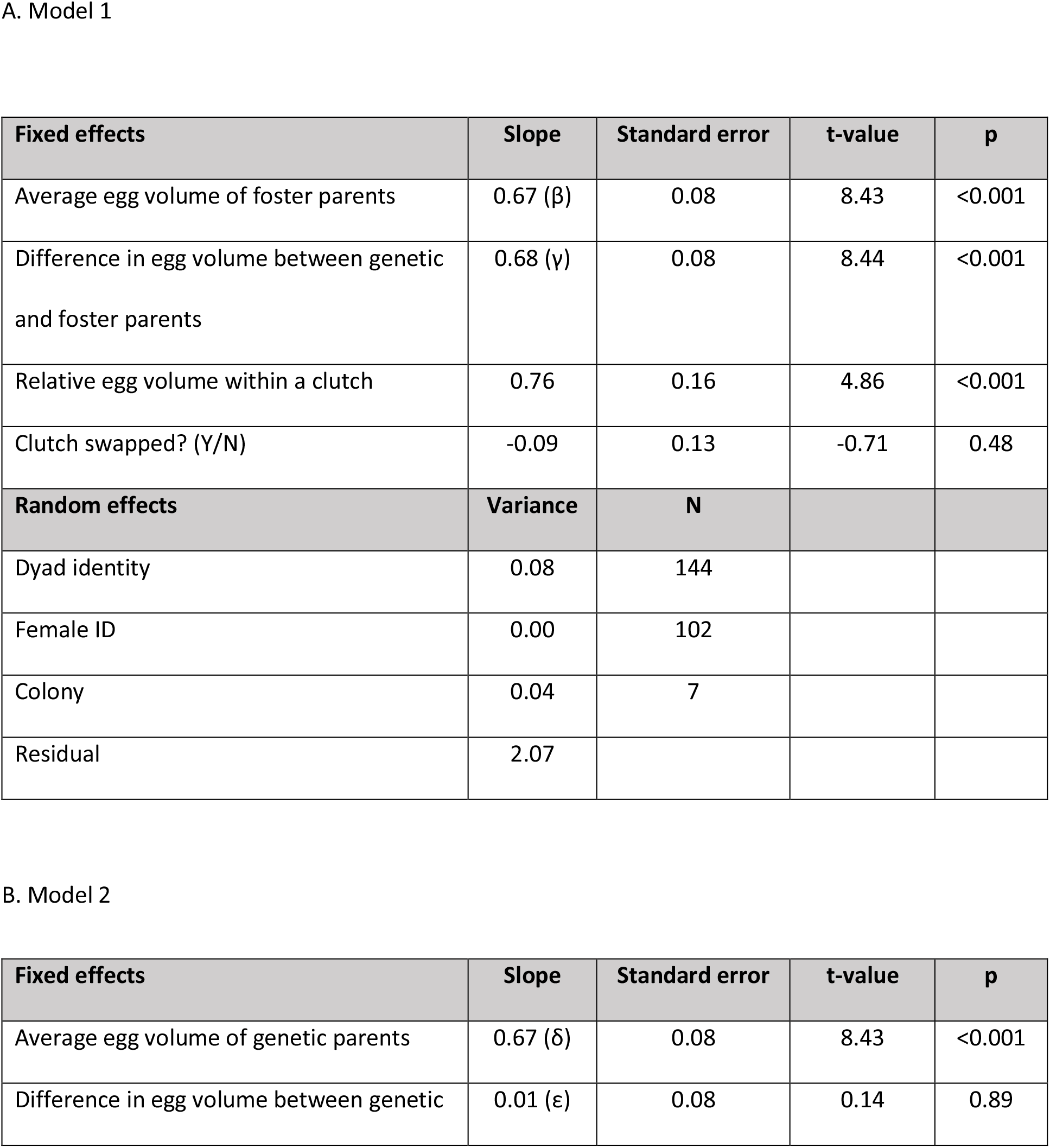
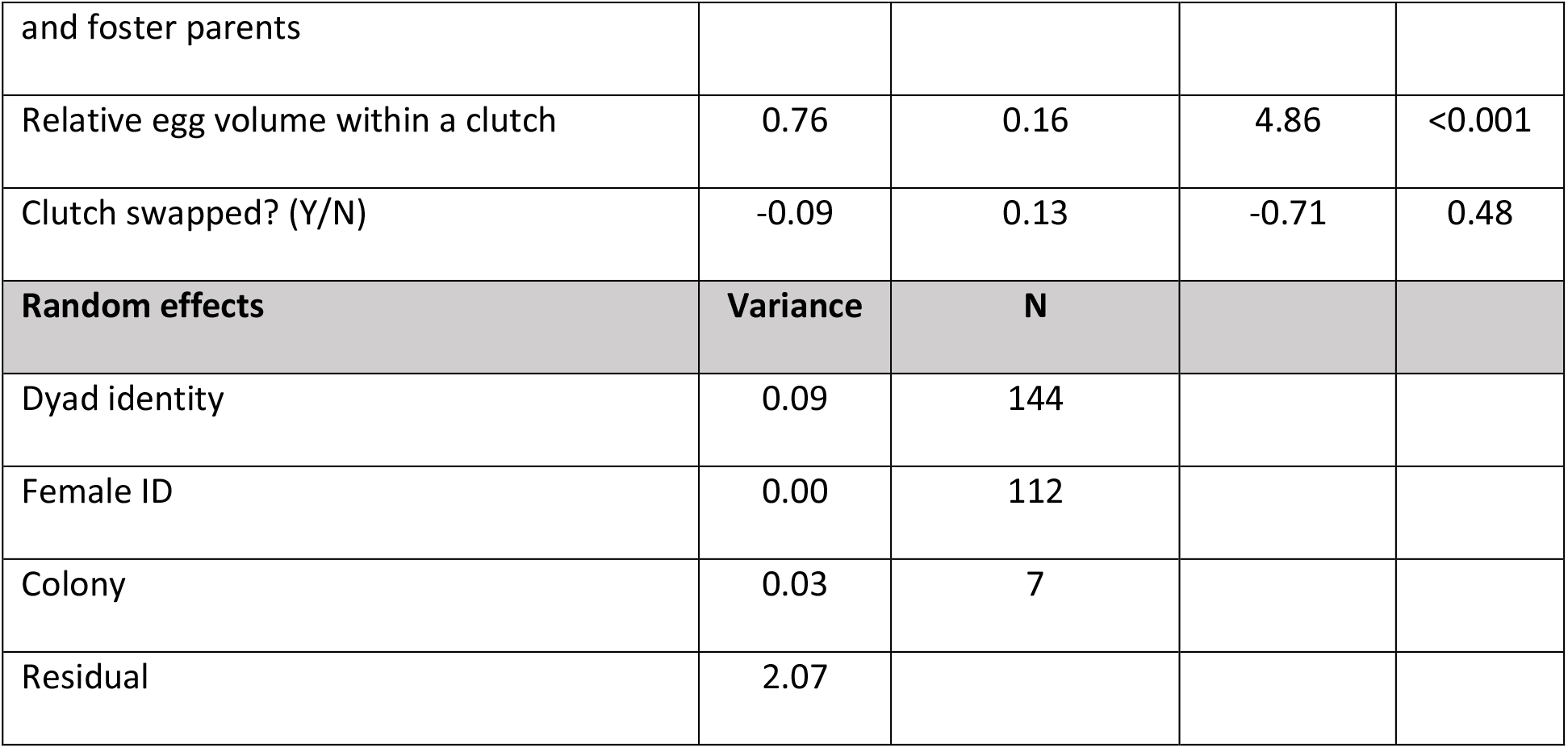
Hatchling mass in relation to egg volume. (a) Effect of egg volume of the foster parents and the difference in egg volume between foster and genetic parents (model 1; marginal R^2^ = 0.16, conditional R^2^ = 0.21). (b) Effect of egg volume of genetic parents and the difference in egg volume between foster and genetic parents (model 2; marginal R^2^ = 0.16, conditional R^2^ = 0.21). Note that female ID includes 11 females whose identity was unknown, and for the purpose of this analysis we assumed them to be 11 uniquely different females.

### Day 5 mass

For full details of model 1 and 2 results, see table 4. Model 1 showed a strong effect of the egg volume of the foster parents (β ± S.E. = 1.82± 0.47 g/cm^3^), but also of the difference in egg volume between the genetic and foster parents (γ ± S.E. = 1.66 ± 0.42 g/cm^3^). These slopes were indistinguishable (Fig. 2), indicating that the association between egg volume and day 5 chick mass can be attributed to egg volume itself (H_1_) and not to parental or environmental quality (H_2_) or a combination of the two (H_3_). This is confirmed by model 2, which showed a strong effect of the egg volume of the genetic parents (δ ± S.E. = 1.82±0.47 g/cm^3^; i.e. the eggs from which the chicks hatched), and no effect of the egg volume difference between the genetic and foster parents (ε ± S.E. = -0.16±0.41 g/cm^3^). Given these coefficients, we estimated that 92% of the association between egg volume and offspring mass can be attributed to egg volume effects, with the remaining 8% attributed to a (non-significant) contribution of other parental effects.

**Table 4:** Nestling day 5 mass in relation to egg volume. (a) Effect of egg volume of the foster parents and the difference in egg volume between genetic and foster parents (model 1; marginal R^2^ = 0.64, conditional R^2^ = 0.73). (b) Effect of egg volume of the genetic parents and the difference in egg volume between genetic and foster parents (model 2; marginal R^2^ = 0.63, conditional R^2^ = 0.73. Note that female ID includes 12 females whose identity was unknown, and for the purpose of this analysis we assumed these were 12 uniquely different females.

A. Model 1.
| Fixed effects | Slope | Standard error | t-value | p |
| --- | --- | --- | --- | --- |
| Average egg volume of foster parents | 1.82 ($\beta$) | 0.47 | 3.41 | <0.001 |
| Difference in egg volume between genetic and foster parents | 1.66 ($\gamma$) | 0.42 | 3.69 | <0.001 |
| Relative egg volume within a clutch | 2.66 | 0.84 | 2.37 | <0.01 |
| Relative hatch date | -13.24 | 0.52 | -24.54 | <0.001 |
| Random effects | Variance | N |  |  |
| Dyad identity | 6.13 | 112 |  |  |
| Female ID | 2.28 | 103 |  |  |
| Colony | 3.61 | 7 |  |  |
| Residual | 34.19 |  |  |  |

B. Model 2.
| Fixed effects | Slope | Standard error | t-value | p-value |
| --- | --- | --- | --- | --- |
| Average egg volume of genetic parents | 1.82 ($\delta$) | 0.47 | 3.84 | <0.001 |
| Difference in egg volume between genetic and foster parents | -0.16 ($\epsilon$) | 0.41 | -0.39 | 0.70 |
| Relative egg volume within a clutch | 2.66 | 0.84 | 3.19 | 0.002 |
| Relative hatch date | -13.24 | 0.52 | -25.28 | <0.001 |
| **Random effects** | **Variance** | **N** |  |  |
| Dyad identity | 6.13 | 112 |  |  |
| Female ID | 2.28 | 103 |  |  |
| Colony | 3.61 | 7 |  |  |
| Residual | 34.19 |  |  |  |

**Figure 2:**
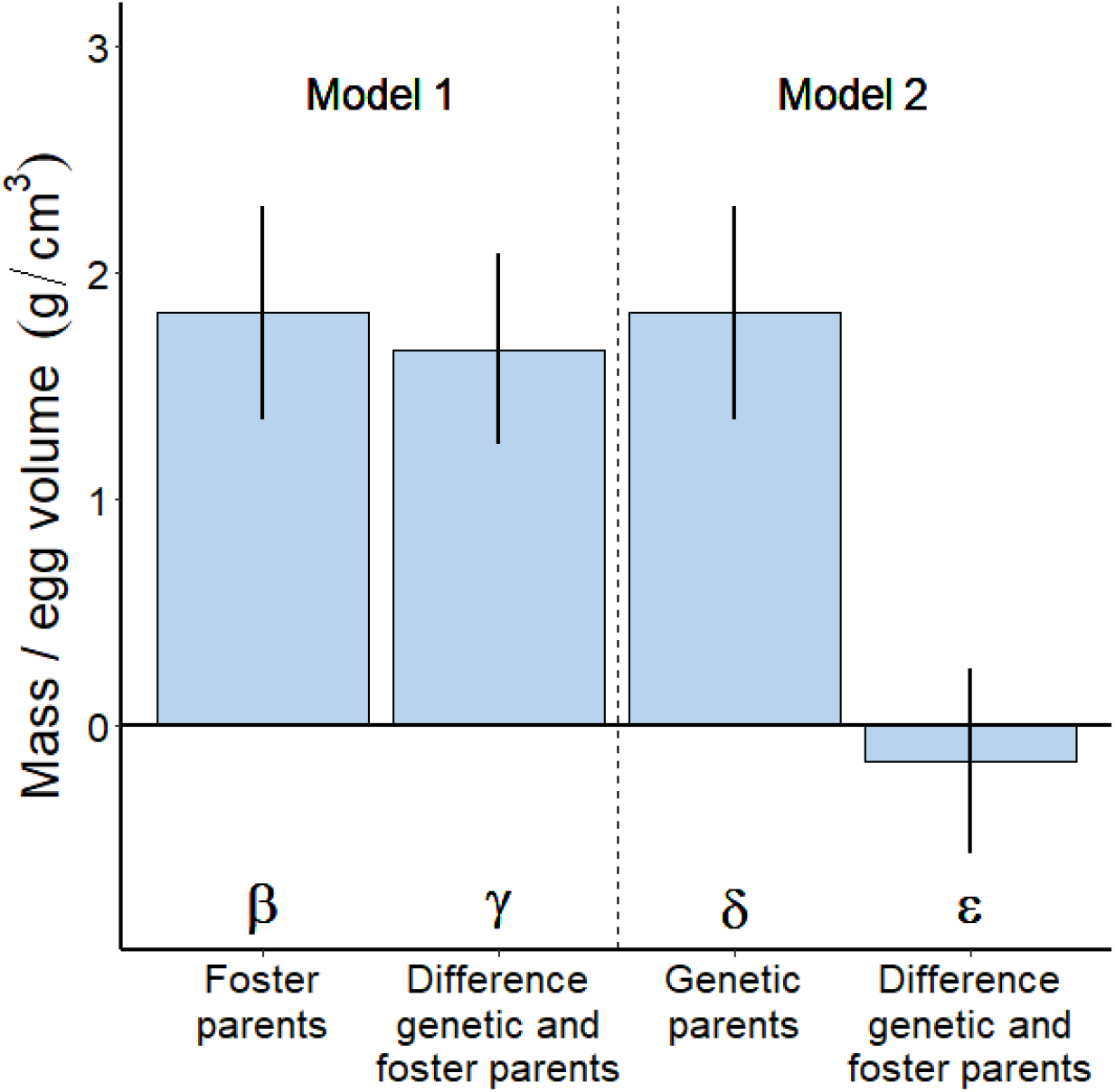
Slope estimates (± S.E.) from models 1 and 2, representing effects on nestling day 5 mass of the average egg volume of the foster parents (β) and the difference in egg volume between genetic and foster parents (γ) in model 1, and the average egg volume of the genetic parent (δ) and the difference in egg volume between genetic and foster parents (ε) in model 2. Note that slope β, γ and δ are indistinguishable, while slope ε is not significantly different from 0. This is in accordance with only direct effects of egg volume on nestling day 5 mass (Hypothesis 1).

**Figure 3:**
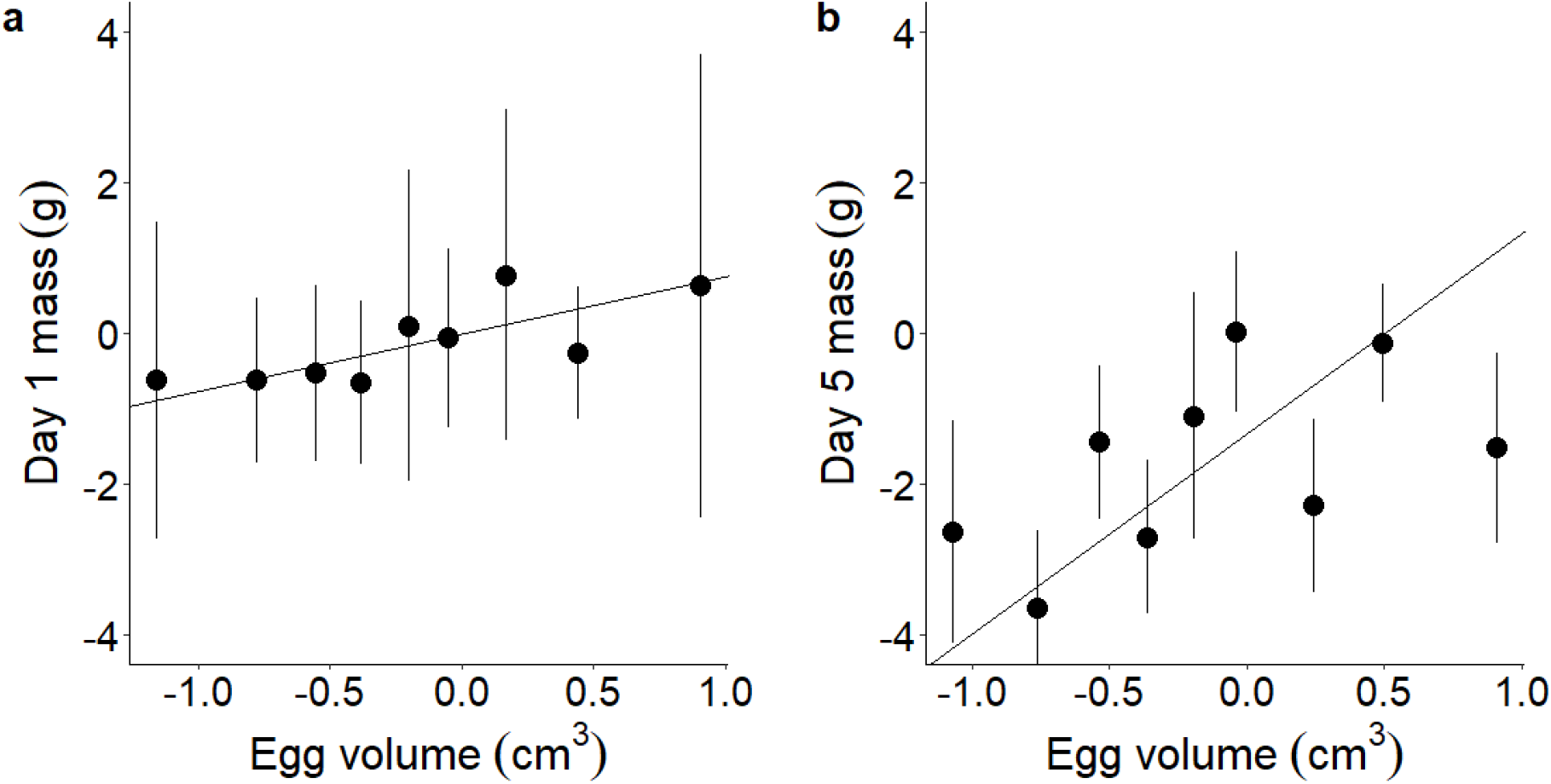
The within clutch effect of egg size on a) hatchling mass and b) nestling day 5 mass. Both egg size and day 5 mass are relative to their clutch average. Only chicks with known egg size were used for this figure. This also causes most data points to be negative, since more late hatched chicks have a known egg size than early hatched chicks. Data points represent averages (± S.E.) of 19 or 15 chicks (a: total number of chicks = 171, from 113 broods, b: total number of chicks = 135, from 91 broods). The lines represent the estimates of the relative egg volume within a clutch from model 1 (a: table 3a, b: table 4a).

The effect of within clutch variation was included in the models by using the difference between individual egg volume and the clutch mean as additional predictor, and also showed a significant effect on day 5 mass (2.7 ± 0.8 g/cm^3^; p<0.01), indicating that within clutch variation in egg volume also affected chick mass (Fig. 3).

We further tested whether the clutch swap affected day 5 mass, but there was no significant effect on day 5 mass of being swapped (coefficient = -0.07 ± 0.83 g, p = 0.93).

### Survival to day 30

To verify the value of day 5 mass as proxy for chick quality, we analysed whether day 5 mass predicted survival until day 30. Brood day 5 mass significantly predicted day 30 survival (Fig.4; slope = 0.04 ± 0.01, p<0.001), as did the individual deviation from the average nestling mass (slope = 0.07 ± 0.007, p<0.001). As expected, brood size manipulation category had a large effect, with higher survival in reduced broods (table 5), and there was no interaction between brood size manipulation and day 5 mass.

**Figure 4:**
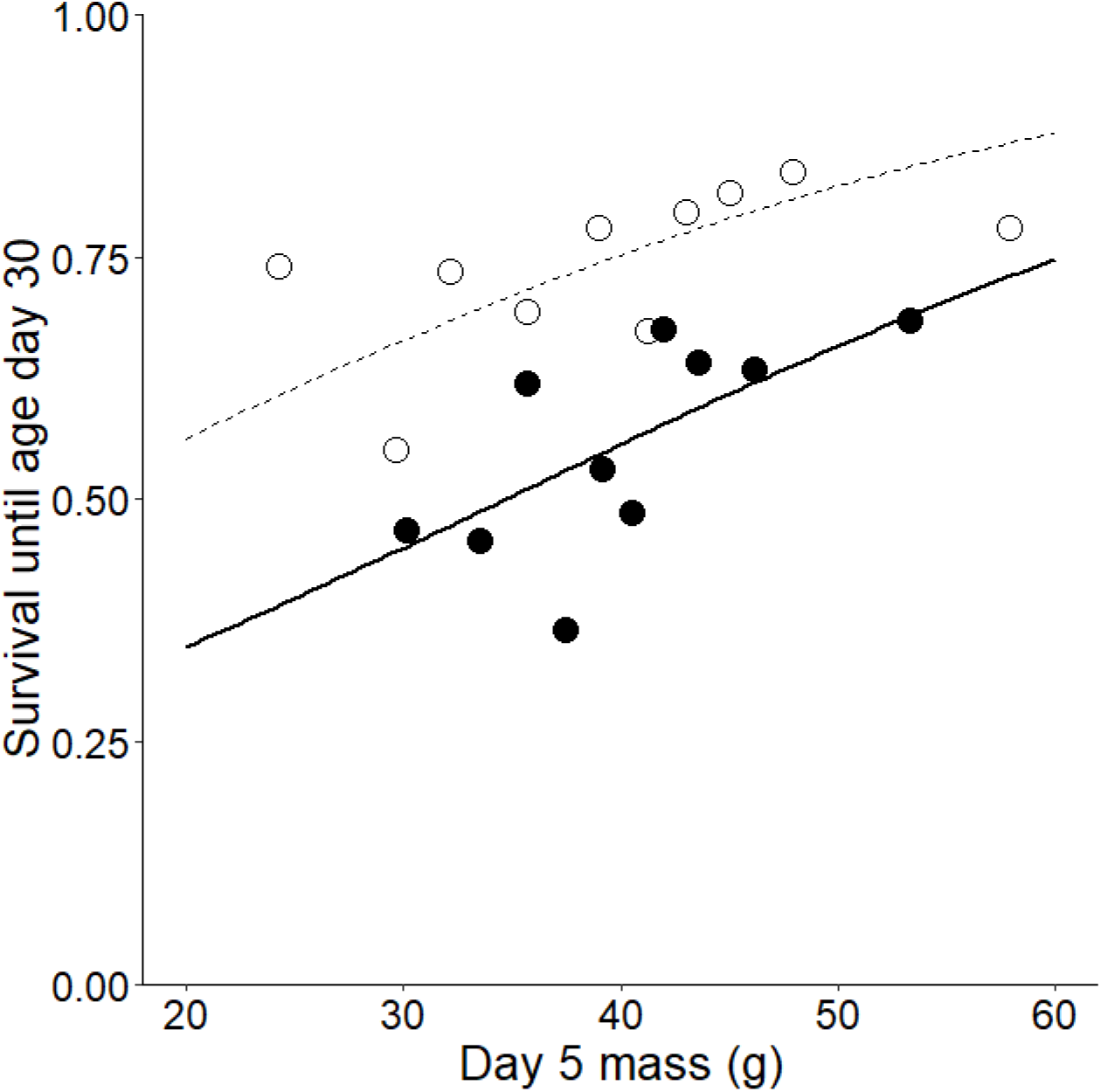
Survival to 30 days as a function of average nestling mass per brood at day 5 in enlarged broods (solid dots and line; n=1423 chicks, from 521 nests) and reduced broods (open dots and dashed line; n = 493 chicks, from 337 nests). The fitted lines are predicted by the model (see table 5). Solid dots are means of 142 (or 143) chicks. Open dots are means of 49 (or 50) chicks.

**Table 5:** Day 5 mass and chick survival until fledging (day 30 of the brood). Because chick survival is binomial, survival was analysed using logistic regression. N = 1914 nestlings in 563 broods. Residual variance is 1 in logistic regressions.

| Fixed Effect | Slope | Standard Error | z-value | p |
| --- | --- | --- | --- | --- |
| Average day 5 mass | 0.04 | 0.01 | 3.63 | <0.001 |
| Relative mass within a clutch | 0.07 | 0.007 | 11.19 | < 0.001 |
| Manipulation Category | -1.20 | 0.16 | -7.66 | <0.001 |
| Random Effect | Variance | N |  |  |
| Female ID | 0.47 | 190 |  |  |
| Year | 2.42 | 13 |  |  |
| Colony | 0.35 | 7 |  |  |

## Discussion

The aim of our study was to investigate whether the association between egg volume and chick mass reported earlier in jackdaws (Verhulst and Salomons 2004) can be attributed to an effect of the egg, an effect of parental care quality or a mixture of both effects. Our results show that this correlation can be attributed to egg volume, with no evidence for additional parental effects to the extent that these are reflected in egg volume. Due to practical constraints, we limited the analysis to day 5 mass, but show that this trait predicts survival up to fledging and therefore conclude that egg volume has an effect on the fitness of the offspring.

Previous cross-foster experiments with the aim to disentangle effects of egg volume and parental quality on offspring performance usually consisted either of swapping complete clutches of large eggs with complete clutches of small eggs, or by randomly swapping complete clutches and adding both the egg volume of the genetic and foster parents in one model to compare which effect explained most of the chick’s quality. We presented an alternative approach to analyse such data, in which hypotheses are tested through the comparison of coefficients of different models (Fig. 1; Table 2). This approach offers two clear advantages. Firstly, in our approach the support for a particular hypothesis is always derived from significant coefficients in the model(s), avoiding inferences made on the basis of the absence of a significant difference, e.g. between pairs having produced small and large eggs. Secondly, the models we constructed allow quantitative inferences on the relative contribution of parents or eggs, through a comparison of the different coefficients. In our study this allowed us to conclude that 92% of the association between egg volume and nestling mass could be attributed to direct effects of egg volume, with the (non-significant) remaining 8% being due to other parental effects. It would be of interest to see these estimates for other populations, allowing a more general assessment of the relative contribution of egg size and other parental effects to associations between egg size and offspring mass. It is further worth noting that application of our approach is not restricted to egg size and body mass, but can be applied generally for the analyses of cross-foster experiments to examine the relative contribution of traits of the parents and the foster parents to the focal phenotypic trait in the offspring.

Egg volume is frequently used as a proxy for parental quality (for example: Bolton 1991, Amundsen et al. 1996, Blomqvist et al. 1997, Monaghan et al. 1998, Hipfner and Gaston 1999, Styrsky et al. 1999, Risch and Rohwer 2000, Bize et al. 2002, Silva et al. 2007, Krist 2009), but is this assumption justified? Since egg volume is positively correlated with chick quality one could argue that parents that lay larger eggs are by definition of a higher quality. This looks like a paradox, since we just concluded that chick quality is not caused by the quality of the parents. However, quality of the parents is multi-dimensional and their ability to feed and take care of their offspring after they are born may be a different component of their quality than their ability to produce large eggs.

An intriguing result emerging from our analysis was that nestling mass at day 5 was 1.7 gram higher for each cm^3^ increase in egg volume, while hatching mass increased only 0.7 g/cm^3^. Apparently, larger eggs not only yield larger hatchlings, but also faster growing nestlings, further amplifying mass differences at hatching. A potential genetic explanation of this observation is that there are pleiotropic genes affecting both egg volume and growth rate. Mechanistically, a way this could be achieved is through the deposition of hormones in the egg, when high concentrations of such hormones stimulates laying large eggs in the female and high growth rate in the hatchling, and assuming that high hormone levels in the female result in high hormone levels in the egg. IGF-1 would be a potential candidate, since it has such diverse effects (Lodjak & Verhulst, 2020). Another potential explanation is that chicks from larger eggs are not only larger at hatching but also further in their development. Since growth rate initially accelerates with age, the difference in mass between further and less developed chicks can then be expected to increase after hatching, explaining the steeper slope of mass on egg volume on day 5 compared to day 1. However, we previously investigated whether egg volume was a predictor of developmental rate as expressed in the incubation time, i.e. the time between laying and hatching of an egg, and found no such effect (Salomons et al. 2006). Sample size has increased considerably in the meantime, but a similar analysis of a much larger data set yielded the same result (Table S2). This suggests that either the incubation time does not indicate developmental rate, with chicks from larger eggs being further in their development at hatching, or that there is another explanation for the observed effect. In either case, this is a puzzle that remains to be resolved.

Although we conclude from our experiment that egg volume affects nestling mass independent of other parental effects, this does not by itself prove that egg volume is causally involved in the observed association. As discussed above, an alternative explanation is that there are genetic effects causing both large eggs and high growth through a mechanism to which the volume of the egg may potentially make a negligible contribution. To demonstrate a causal effect of egg volume per se, it would be necessary to manipulate the size of the egg directly. This has been achieved through surgical ablation of eggs in lizards (Sinervo and Licht 1991), but performing a comparable manipulation in wild birds will be challenging.

Our clutch swaps were on average performed seven days after clutch completion, to be sure the clutch was really completed – jackdaws sometimes have a laying stop for one or multiple days and we checked nests every three days during laying. In these 7 days the genetic mother could still have had some effects on the chicks via incubation. However, we did not see an effect of egg volume on incubation time (Table S1), so we therefore do not expect that incubation behaviour of the mother affects egg sizes differently or that the delay in clutch swap changes our results.

The finding that larger eggs have a positive effect on nestling mass raises the question why there is still variation in egg size, and what causes this variation. Part of this variation is likely to have a genetic origin, because heritability of egg size is often considerable (Larsson and Forslund 1992, Potti 1999), but this remains to be estimated for our population. We previously reported that egg size was positively correlated with size-adjusted mass of females shortly after hatching (Verhulst and Salomons 2004), suggesting that variation in egg size at least partly reflects variation in the resources available for the female to invest in reproduction. Further considerations that could lead to egg size being smaller than the maximum are that there may be a trade-off between the quality and quantity of eggs (Sinervo and Licht 1991, Hendry et al. 2001, Fischer et al. 2011) and a trade-off between investment in the clutch and subsequent survival and reproduction (Nager et al. 2001). For example, Monaghan et al. (1998) reported that when parents were experimentally induced to lay an extra egg, they were less successful at rearing (foster) chicks. Indeed, females in our study population restrained their investment in eggs, as illustrated by the finding that they increased the size of their eggs when their life expectancy was experimentally shortened (Boonekamp et al 2020). Thus, egg size is a complex trait and attaining a complete understanding of the causes of its variation remains a challenge.

## Acknowledgements

We would like to thank Martijn Salomons, the late Cor Dijkstra, all students that helped collecting the data over the years, and reviewers of an earlier version of this paper for helping us to clarify our approach.

## Ethics statement

This work was carried out under license numbers (4071, 5871, 6832-A & 6832-B).

## Data availability

Data will be available at the University of Groningen Dataverse once this manuscript is published, using this link: https://doi.org/10.34894/II4NMB.

## Appendix 1

Here we show how we derived the predictions for the model coefficients given the different hypotheses.

### Hypothesis 1 – Only egg effects

Model 3

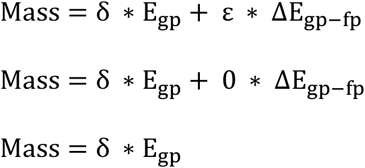

Model 2

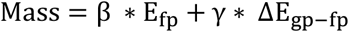

Model 3 can be rewritten:

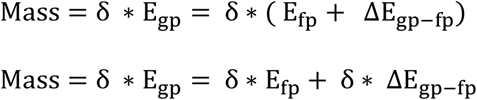

Model 2 and model 3 are always equal, thus

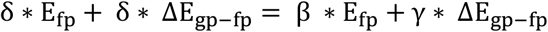

Which holds true when β = γ = δ (and ε = 0 as there are only egg effects).

### Hypothesis 2 – Only parental effects

Model 2

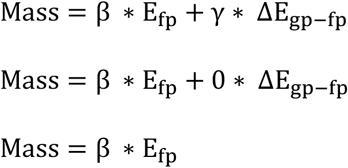

Model 3

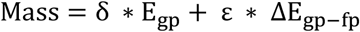

Model 2 can be rewritten:

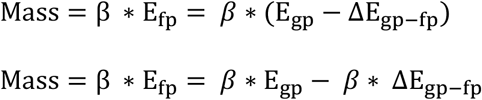

Model 2 and model 3 are always equal, thus

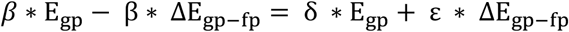

Which holds true when β = δ = -ε (and γ = 0 as there are only parental effects)

