## Supplementary Figure S1 and Table S1 for "Egg size effects on nestling mass in Jackdaws *Corvus monedula*: a cross-foster experiment"

### Supplementary information

This supplement contains two appendices:

Appendix A: Supplementary figure S1.

Appendix B: Supplementary table S2.

**Appendix A: Supplementary figure**

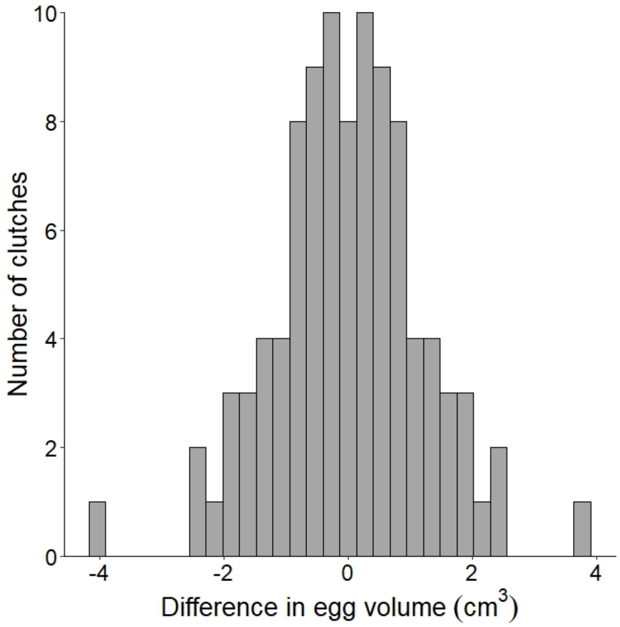


Figure S1: A histogram showing the difference in average egg volume between the swapped clutches. The clutches were swapped matched for clutch size and laying date, irrespective of egg volume. The average egg volume of these clutches was 10.81 $cm^3$.

**Appendix B: Supplementary table**

Table S1: Effects of egg volume and clutch size on incubation time in days. N = 682 eggs in 195 clutches. Eggs were kept in an incubator after clutch completion until hatching. For detailed information about the methods see (Salomons et al. 2006).

| | **Fixed Effect** | **Slope** | **Standard Error** | **p-value** |
|---|---|---|---|---|
| **Covariates** | Egg volume | -0.047 | 0.036 | 0.19 |
| **Factors** | | | | |
| Laying Order | $1^{st}$ egg – $1.5^{th}$ egg | -0.26 | 0.12 | 0.033 |
| | $1^{st}$ egg – $2^{nd}$ egg | -0.97 | 0.07 | <0.001 |
| | $1^{st}$ egg – $2.5^{th}$ egg | -0.94 | 0.11 | <0.001 |
| | $1^{st}$ egg – $3^{rd}$ egg | -1.73 | 0.07 | <0.001 |
| | $1^{st}$ egg – $3.5^{th}$ egg | -1.64 | 0.13 | <0.001 |
| | $1^{st}$ egg – $4^{th}$ egg | -2.27 | 0.08 | <0.001 |
| | $1^{st}$ egg – $4.5^{th}$ egg | -2.05 | 0.14 | <0.001 |
| | $1^{st}$ egg – $5^{th}$ egg | -2.81 | 0.09 | <0.001 |
| | $1^{st}$ egg – $6^{th}$ egg | -3.05 | 0.24 | <0.001 |
| | $1^{st}$ egg – $8^{th}$ egg | -3.55 | 0.57 | <0.001 |
| Clutch size | 1 egg – 2 eggs | -1.02 | 1.15 | 0.38 |
| | 1 egg – 3 eggs | -0.21 | 0.83 | 0.80 |
| | 1 egg – 4 eggs | 0.04 | 0.82 | 0.97 |

| | | | | |
|---|---|---|---|---|
| | 1 egg – 5 eggs | 0.28 | 0.82 | 0.73 |
| | 1 egg – 6 eggs | 0.87 | 0.85 | 0.31 |
| | **Random effect** | **Variance** | **Number of groups** | |
| | NestID | 0.378 | 195 | |
| | Year | 0.015 | 11 | |
| | Residual | 0.272 | | |
